# AI-driven high-throughput droplet screening of cell-free gene expression

**DOI:** 10.1101/2024.07.04.602084

**Authors:** Jiawei Zhu, Yaru Meng, Wenli Gao, Shuo Yang, Wenjie Zhu, Xiangyang Ji, Xuanpei Zhai, Wan-Qiu Liu, Yuan Luo, Shengjie Ling, Jian Li, Yifan Liu

**Author notes:** Corresponding author. (S.L.), (J.L.), (Y.L.). These authors contribute equally to this work (Jiawei Zhu, Yaru Meng, Wenli Gao).

## Abstract

Cell-free gene expression (CFE) systems enable transcription and translation using crude cellular extracts, offering a versatile platform for synthetic biology by eliminating the need to maintain living cells. This allows direct manipulation of molecular components and the focused synthesis of specific products. However, the optimization of CFE systems is constrained by cumbersome composition, high costs, and limited yields due to numerous additional components required to maintain biocatalytic efficiency. While optimizing such complicated systems is daunting for existing high-throughput screening means, we introduce DropAI, a droplet-based, AI-driven screening strategy designed to optimize CFE systems with high throughput and economic efficiency. DropAI employs microfluidics to generate picoliter reactors and utilizes a fluorescent color-based coding-decoding system to address and screen a vast array of additive combinations. The in-droplet screening is complemented by in silico optimization, where experimental results train a machine-learning model to estimate the contribution of the components and predict high-yield combinations, which are then validated in vitro. Applying DropAI to an *Escherichia coli*-based CFE system, we simplified a set of 12 additives to only 3 essential components. Through further optimization, we achieved a 2.1-fold cost reduction and a 1.9-fold increase in yield for the expression of superfolder green fluorescent protein (sfGFP). This optimized formulation was further validated across 12 different proteins. Notably, the established *E. coli* model is successfully adapted to a *Bacillus subtilis*-based system through transfer learning, leading to doubled yield through prediction. DropAI thus offers a generalizable and scalable method for optimizing CFE systems, enhancing their potential for biochemical engineering and biomanufacturing applications.

## 1. Introduction

Cell-free gene expression (CFE) involves the in vitro activation of transcription and translation using crude cellular extracts instead of intact cells^1^. Compared to in vivo gene expression, CFE offers a more flexible, sustainable, and rapid approach to gene expression. First, by eliminating the necessity of sustaining cellular life, it allows researchers to directly manipulate the molecular environment of the system. This includes the addition of non-native substrates, purified proteins or RNAs, and recombinant DNA templates. Furthermore, CFE circumvents mechanisms essential for cell survival, bypasses limitations on molecular transport across the cell wall, and enables a focused investigation on specific genetic networks or the biosynthesis of a single product. Moreover, CFE can accelerate design-build-test cycles in biotechnological applications by significantly reducing the timeline of cloning steps and gene expression. Given these advantages, there has been a resurgence of scientific interest in cell-free biotechnology, leading to the development of novel CFE platforms for various compelling applications^1,2^, such as industrially important chemical production^3-6^, metabolic pathway prototyping^7^, natural product biosynthesis^5^, clinical therapeutics^8,9^, and diagnostics^10^.

However, standardized procedures for optimizing each newly developed CFE system still remain elusive, which are often dependent on empirical approaches. Moreover, current CFE systems are constrained by the complicated formulations and the challenge of balancing cost and yield^1^, which severely limits their utility in an expanded repertoire of applications. For example, a typical bacteria extract-based CFE system requires around 40 additional components, besides the crude cell extract, to maintain a reasonable level of biocatalytic efficiency. Over half of the cost of a standard bench-scale CFE formulation arises from the expensive energy substrates and multiple additives used to enhance biosynthesis yield^1^. While some of these additives, such as dNTPs and nucleic acids, are essential, many others can be optimized or minimized. Addressing these challenges calls for comprehensive screening of the CFE system to develop a streamlined formulation with cost-effective, sustainable energy sources and minimal essential additives at optimized concentrations. Such screening not only has the potential to reduce costs but also to provide new insights into CFE as an integrated biochemical system^11^.

High-throughput screening (HTS) has found widespread use in drug discovery^12-16^ and biochemical system optimization^17^. Still, the adoption of HTS in CFE optimization has been hindered by its complexity, compound consumption, and cost^1^. A typical CFE formulation includes non-native energy sources, gene templates, and various transcription-translation factors (e.g., coenzyme A, NAD, and exogenous tRNAs). Screening these additives can easily generate thousands of combinations. For example, considering five candidate energy source molecules (choose one out of five) and ten different transcription-translation factors (only consider their presence or absence), a primary screening process must evaluate 5,120 combinations (5×2^10^). Constructing a screening pool at this scale requires approximately 60,000 liquid handling steps and 54×96-well plates (without replication). Given the logistical complexity, conventional pipette-based liquid handling techniques are often inefficient or unfeasible. The screening scale can expand exponentially when introducing concentration gradients of the additives. Additionally, CFE formulations vary with the choice of chassis cells (e.g., *Escherichia coli*^18^, *Streptomyces*^19^, and yeast^20^). Consequently, changing the chassis cell extract necessitates repeating the entire screening process to optimize the CFE system. Given these challenges, adequately screening CFE systems with existing HTS methods is impractical.

Here, we report a droplet-based and AI-guided combinatorial screening strategy (DropAI) designed to optimize CFE systems with high throughput and economic efficiency. A key feature of DropAI is its ability to construct massive combinations in picoliter reactors (80-μm droplets, ∼250 pL) and use a fluorescent color-based coding-decoding approach to trace the composition of each combination (Fig. 1a). This is achieved through microfluidics, where one carrier droplet merges with four satellite droplets to form a complete screening unit. The carrier droplet contains the CFE mixture, while each satellite droplet randomly samples a unique set of CFE components. These component sets are labeled with distinct fluorescent colors, and each component within a set is associated with a unique fluorescence intensity. Consequently, the merged droplets are encoded by fluorescent colors and intensities, termed FluoreCode, to identify the contained combinations. The FluoreCode is read in parallel using multi-channel droplet imaging. With our current optical settings, we can resolve nine intensity levels within one color, enabling the satellite droplets to encode 729 (9^3^) combinations. If the carrier droplets are also color-coded, the theoretical combinatorial space expands to 6,561 (9^4^). In microfluidics, combinations are created at a rate of approximately 1,000,000/hour. For a screening scale of 500 combinations with 100 replicates each (totaling 50,000 droplets), it takes about three minutes to construct the entire pool (excluding a setup time of around 30 minutes), consuming approximately 12.5 μL of reagents.

**Figure 1.**
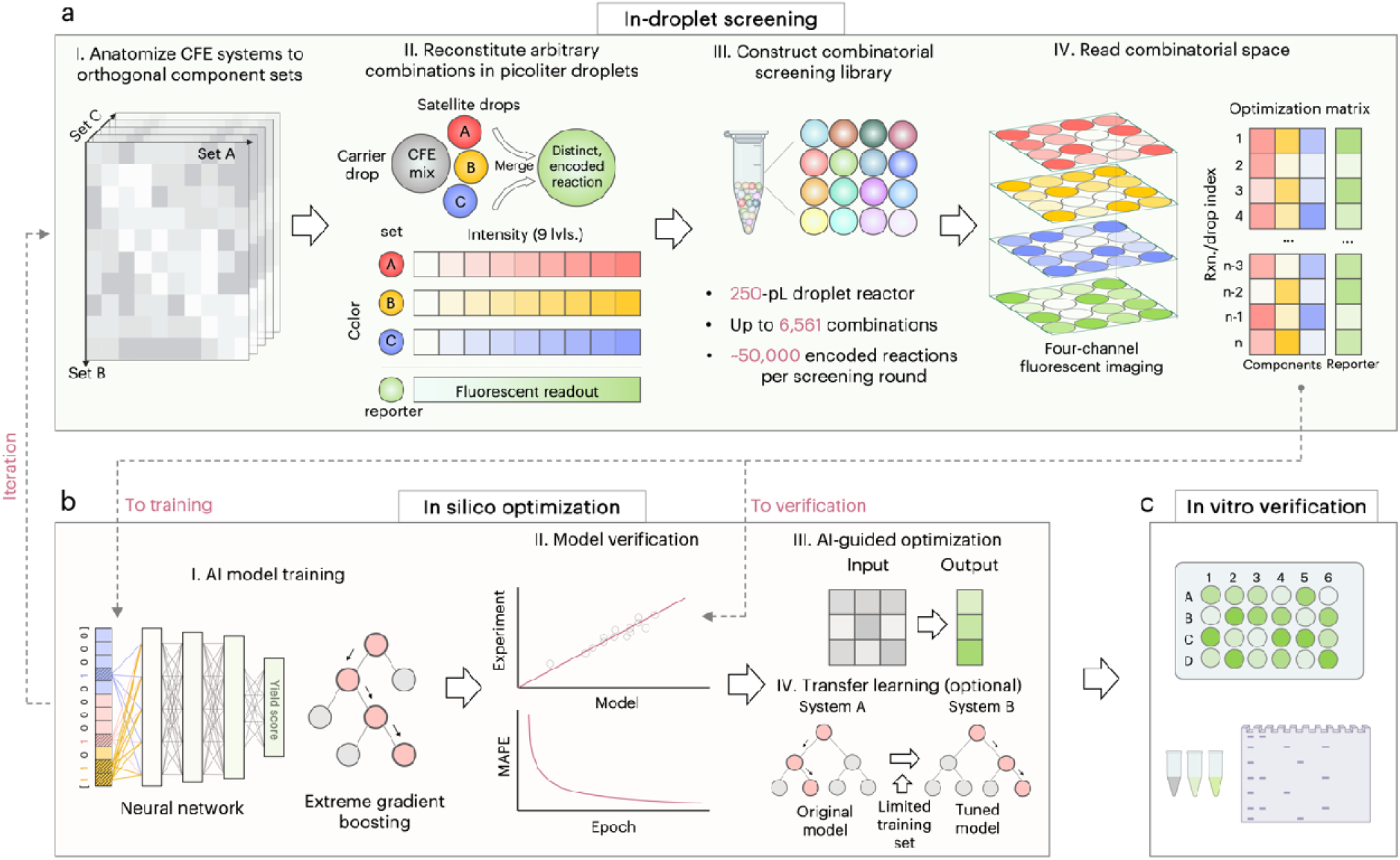
DropAI Workflow. DropAI employs a hybrid strategy of (a) in-droplet screening and (b) in silico optimization. (a) In-droplet screening: Massively parallel experimental data are generated using picoliter droplet reactors. Each droplet contains a unique combination of CFE components encoded by fluorescent markers. Multichannel droplet imaging allows for parallel reading of component combinations and resulting CFE yields (sfGFP fluorescence). (b) In silico optimization: Experimental data are used to develop AI models, which extend predictions beyond experimental scales to enhance screening capacity. Iterations of the screening process may be conducted for further optimization. (c) Final verification: The optimized screening results are validated in vitro.

After the in-droplet screening, DropAI proceeds with in silico optimization (Fig. 1b). Experimental results obtained from the droplet assays are used to train a machine learning model. The established model explores conditions beyond the experimental scale to predict high-yield combinations, which are then verified in vitro (Fig. 1c). In this study, we first applied DropAI to optimize an *E. coli*-based CFE system. Using in-droplet screening, we sampled combinations of 12 additives across transcription, translation, and ATP regeneration phases of CFE. We then built a machine learning model based on the experimental data to predict the contribution of each additive to CFE yield. Based on these predictions, we selected only three essential additives and optimized their concentrations. The final simplified and optimized CFE formula was tested by expressing superfolder green fluorescent protein (sfGFP), resulting in a 2.1-fold decrease in unit cost and a 1.9-fold increase in yield compared to the original formula. We further applied the optimized formula to express 12 different proteins ranging from 27 to 370 kDa, with 10 out of 12 proteins exhibiting maintained or increased yields. Furthermore, we demonstrated that the established *E. coli* model could be tuned and applied to optimize a *Bacillus subtilis*-based CFE system through transfer learning^21^.

This approach significantly reduced the effort required to optimize CFE systems across different chassis cells. The transfer model achieved high prediction accuracy with a limited number of new experiments (e.g., 27 and 36 combinatorial sets). The predicted high-yield formulation was verified in vitro by expressing sfGFP (resulting in a two-fold increase in yield) and short antimicrobial peptides.

## 2. Results

### 2.1. Microfluidic construction of combinatorial droplet library

DropAI relies on high-throughput microfluidics to generate massive combinatorial libraries for in-droplet screening. To realize this concept, we developed a microfluidic device capable of generating arbitrary four-droplet combinations at approximately 300 Hz (Fig. 2a and Supplementary Fig. 1a). In this device, a pool of carrier droplets (80 μm in diameter) is loaded upstream in a microchannel. The carrier droplets flow downstream sequentially, each meeting three satellite droplets (36 μm in diameter) before reaching a micro-teeth structure designed for droplet merging (Fig. 2b and Supplementary Figs. 1b&c). This process is achieved by synchronizing the frequency of droplets from different pools, resulting in an overall efficiency of approximately 80%. Each droplet is encoded with a unique fluorescent color and intensity, creating a final merged droplet with a 4-digit nonary FluoreCode (Fig. 2c). To evaluate the coding capacity, we generated a library of 6,561 (9^4^) combinations using fluorescent dye solutions (Fig. 2d). The droplets were imaged under multiple channels (Fig. 2e and Supplementary Fig. 1d) to extract the fluorescence of each droplet (n = 206,634). We confirmed that the intensity profile of each color exhibited nine major peaks (Supplementary Fig. 1e). We then processed the intensity data using a multi-band-pass filter program to eliminate low-quality marginal data points. The remaining data sets (n = 153,302) were dimensionally reduced and clustered using t-distributed stochastic neighbor embedding (t-SNE), as plotted in Fig. 2f. The t-SNE analysis recognized 6,527 distinct clusters, recovering 99.5% of the theoretical combinatorial space.

**Figure 2.**
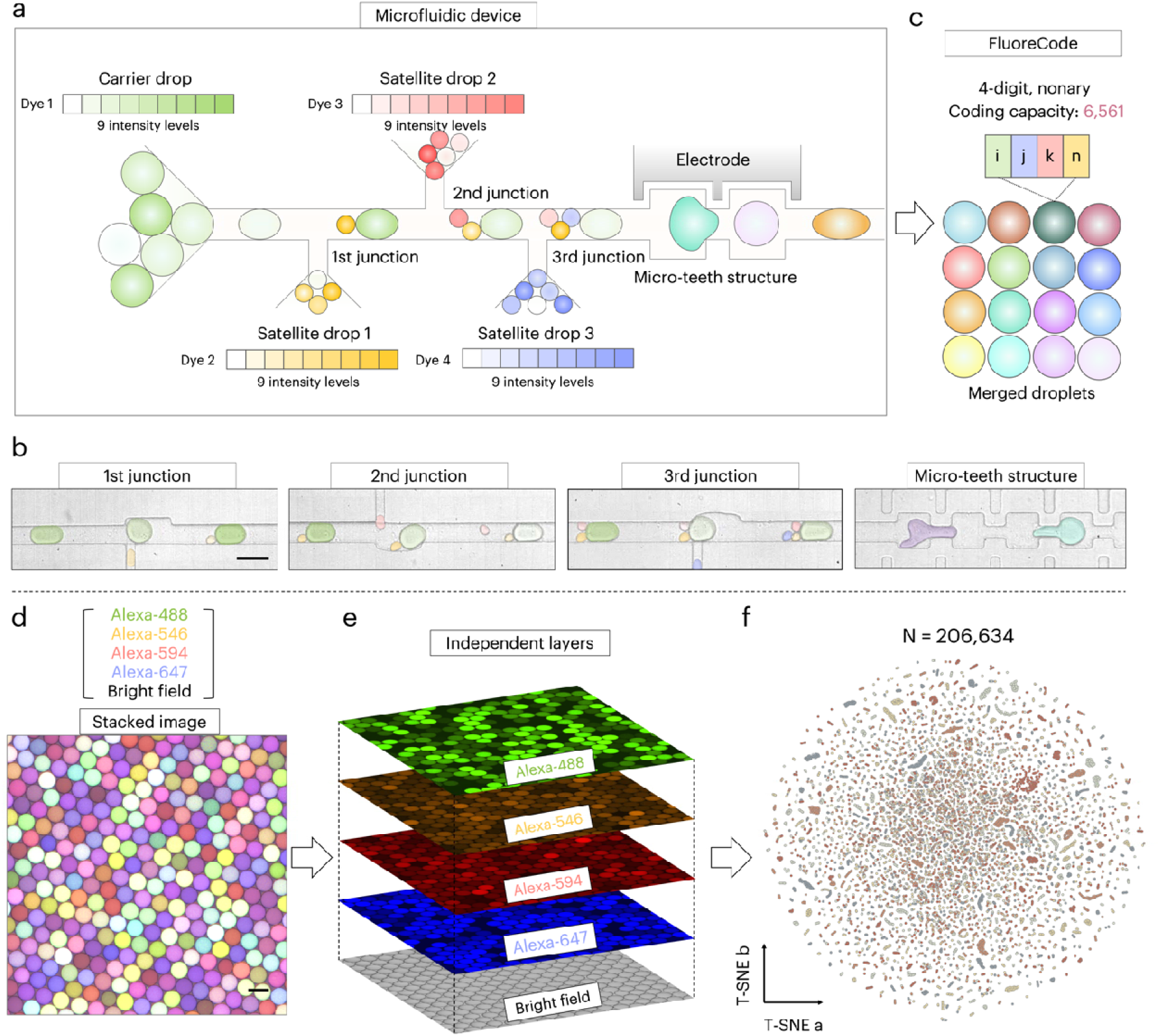
Microfluidics and coding/decoding in DropAI. (a) DropAI generates massive combinations in droplet reactors with a multi-droplet microfluidic merger. The device loads a pool of carrier droplets into a microchannel at regular intervals. Each carrier droplet meets 3 satellite droplets at downstream junctions, which are all merged under an electrical field at the micro-teeth structure. (b) Micrographs depict the droplet pairing and merging process. (c) As every droplet is coded by fluorescent color and intensity, the merged droplets are rendered a 4-color barcode (FluoreCode) indicating the exact combinations. The microfluidic device generates the combinatorial pool at ∼300 Hz, and the total coding capacity is 6,561 (9^4^). (d) A stacked micrograph displaying the as-built combinatorial pool. (e) For decoding, the merged droplets are imaged under 4 fluorescent channels to extract the FluoreCode of each droplet. The droplets are then clustered based on the FluoreCode information. (f) A t-distributed stochastic neighbor embedding (T-SNE) projection of 206,634 droplets depicting 6,527 clusters, ∼99.5% of the entire combinatorial space. Scale bars: 100 μm.

### 2.2. Validation of CFE in DropAI

Having established the microfluidics, we next examined the compatibility of CFE in the DropAI workflow. We first asked whether CFE could be carried out in the picoliter droplet reactors. To investigate this, we emulsified an *E. coli* lysate-based CFE assay into 80-μm droplets using a microfluidic flow-focusing chip (Supplementary Fig. 1a), with fluorinated oil and a biocompatible polyethylene glycol-perfluoro polyether (PEG-PFPE) surfactant (Fig. 2a). The assay was designed to express a reporter protein, sfGFP. After incubation, the droplets exhibited notable fluorescence, indicating successful expression of sfGFP (Supplementary Fig. 2a). However, the emulsions were not mechanically stable, as large collapsed droplets were observed (Supplementary Fig. 2b). To stabilize the emulsions, we added Poloxamer 188 (P-188), a non-ionic triblock-copolymer surfactant, and Polyethylene glycol 6000 (PEG-6000), a biocompatible crowding reagent, to the aqueous phase (CFE mix). These polymers have been verified as effective stabilizers for droplet-based biochemical assays. Indeed, after adding P-188 and PEG-6000, the droplets remained intact throughout the incubation process (Supplementary Fig. 2c). Next, we extracted the aqueous phase from the emulsions and measured the sfGFP fluorescence. The fluorescence level was comparable to that of a parallel bulk CFE assay (standard 15 μL reactions in 1.5 mL test tubes), suggesting that CFE efficiency is identical in both systems (Supplementary Fig. 2d).

We then investigated whether CFE is compatible with droplet fluorescence barcoding. To test this, we first conducted droplet-based CFE in the presence of dye molecules (AlexaFluor 546) and found that adding fluorescent dye did not affect protein expression (Supplementary Fig. 2e). In addition, we confirmed that the dye molecules remained in the aqueous phase without diffusing into the surrounding oil for up to 24 hours (Supplementary Fig. 2f). Thus, the fluorescence coding resolution would not be compromised by dye molecules transporting across droplets during the CFE incubation (4 hours). Next, we encoded the presence of Mg^2+^, an essential ion in CFE, to test the fluorescence coding and decoding. As depicted in Fig. 3b, the droplets formed two distinct clusters, and the occurrence of sfGFP fluorescence correlated with the encoded presence of Mg^2+^. Encouraged by this result, we further encoded a five-grade concentration gradient of Mg^2+^. As shown in Fig. 3c, five distinct clusters of droplets were clearly resolved. The sfGFP levels in these clusters indicated that sfGFP expression peaked at 12 mM Mg^2+^, which was confirmed by identical experiments performed in bulk (Fig. 3c inset). Overall, these results validate that DropAI is a feasible and efficient approach for optimizing CFE systems.

**Figure 3.**
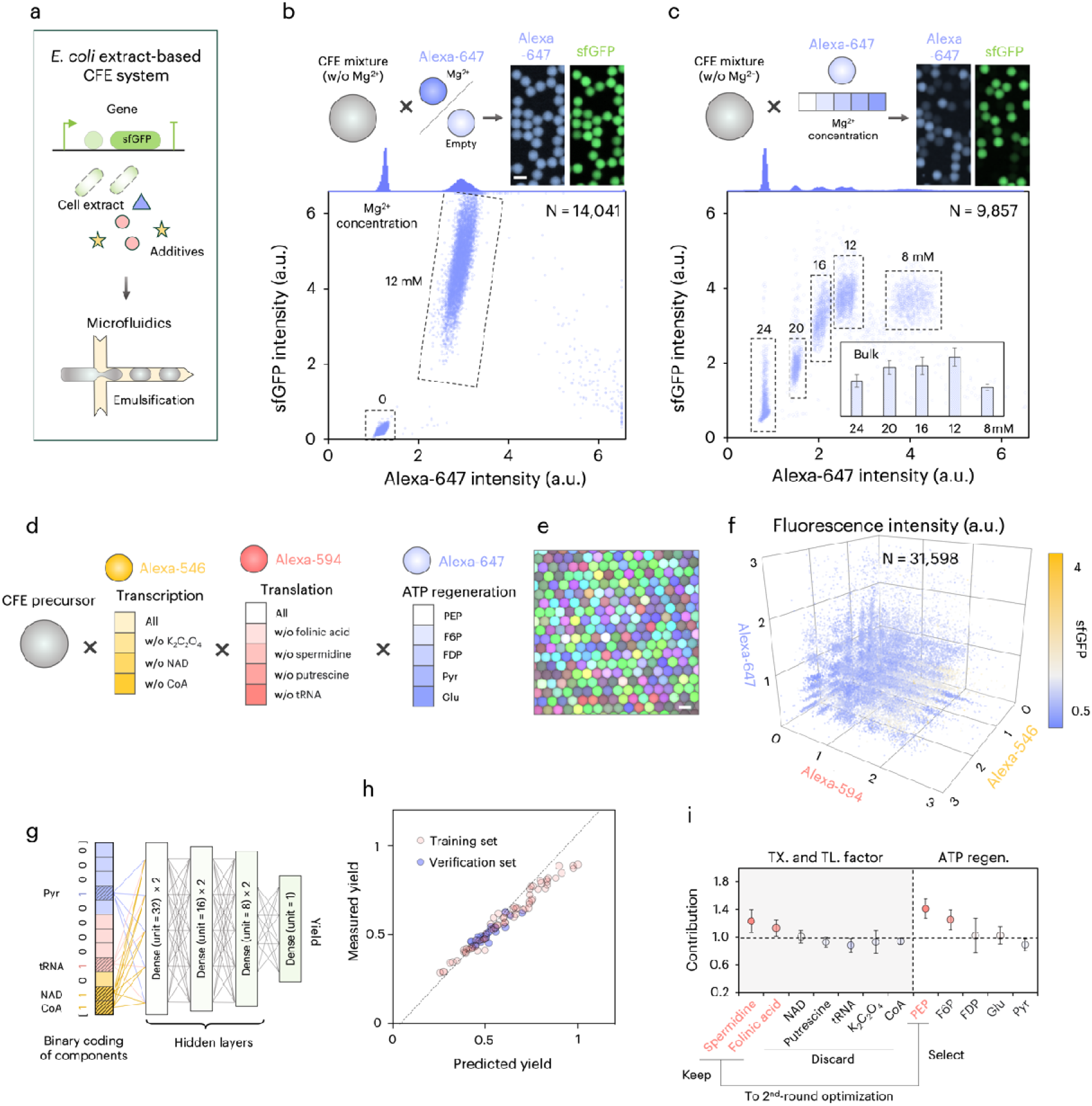
Primary optimization of an *E. coli* extract-based CFE system. (a) Illustration of the CFE mixture composition and emulsification process. (b&c) Preliminary experiments encoding the (b) presence and (c) concentrations of Mg^2+^. The upper panels display the encoding process and post-CFE droplets under the encoding dye (Alexa-647) and sfGFP channels. The scatter plot on the bottom reveals the fluorescence intensity profile of the droplets. The inset column plot in (c) depicts a parallel experiment performed in bulk (15 μL in-tube). (d-f) In-droplet screening of 12 CFE components. (d) The 3-color coding strategy. (e) A representative micrograph of the encoding droplets (fake colors are used for display purposes). (f) A 3D scatter plot showing the fluorescence intensity profile of the droplets. The color of the dots indicates the sfGFP intensity. (g-h) In silico optimization. (g) A 7-layer neural network model used in the optimization. The model interprets the presence of components as binary values and outputs the contribution score of each component. (h) Comparison of the sfGFP yield (fluorescence intensity) obtained from the droplets vs. the model predictions. (i) The in silico estimated contribution of the 12 components to the protein yield. Spermidine, Folinic acid, and PEP are selected for the second-round optimization. Scale bars: 100 μm.

### 2.3. Combinatorial screening of *E. coli*-based CFE

After proving the concept, we proceeded to use DropAI to optimize a CFE system derived from *E. coli*, which is well-developed with standardized formulations and widely used in various laboratories^18,22^. Our goal was to achieve a simplified formulation by identifying key additives essential for protein expression and discarding non-essential ones. Additionally, we aimed to improve the overall yield by optimizing the concentrations of the identified key additives.

We conducted two rounds of DropAI screening using the *E. coli*-based PANOx-SP CFE system, following standardized protocols^22^. The first round aimed to determine the minimal set of essential additives, and the second round aimed to optimize their concentrations. In the first round, we started with 12 widely-used components from the PANOx-SP system, including 7 major additives for transcription-translation and 5 energy suppliers for ATP regeneration. We used the widely suggested concentrations of the components^22^ (Supplementary Table 1) and only encoded their presence. Note that each combination was set to have no more than one energy substrate. Thus, these components constituted a combinatorial space of 768 [2^7^×(5+1)]. For the in-droplet screening, we applied 3 fluorescent colors to encode a subset of the combinatorial space (100 combinations) and used sfGFP yield (fluorescence intensity) as the output (Fig. 3d). As the carrier droplets were not coded, we used a simplified microfluidic device (Supplementary Fig. 1a) where the carrier drops were produced directly on-chip.

The in-droplet screening (Fig. 3e) generated 31,598 droplet-based CFE data points (Fig. 3f), which were binned into 100 clusters according to the FluoreCode and fed into a neural network model (Fig. 3g). The model consisted of 7 fully connected layers and interpreted the presence of each additive as binary values, with normalized sfGFP fluorescence as the output (Supplementary Fig. 3a). Among the input data, 70 combinatorial sets were used to train the model and 30 sets to test its prediction accuracy. During the training process, the mean absolute percentage error (MAPE) loss function converged to ∼6.9% after 1,000 training epochs without obvious overfitting (Supplementary Fig. 3b). After training, we obtained an R² of ∼0.86 and a linear fit with an intercept of -0.08 and a slope of 1.18 (Fig. 3h). These results verified the accuracy of the machine learning model.

The established model scanned through the entire combinatorial space and scored the contribution of the 12 compounds to protein yield by calculating the average ratio between the yields with and without each compound (Fig. 3i and Supplementary Fig. 3c). Among the transcription-translation factors, spermidine and folinic acid exhibited scores greater than 1.1, suggesting that their presence alone can increase the yield of the CFE system by more than 10%. However, NAD, putrescine, tRNA, K_2_C_2_O_4_, and CoA were deemed non-essential for protein yield. Among the energy sources, phosphorylated energy substrates [phosphoenolpyruvate (PEP), fructose 6-phosphate (F6P), and fructose-1,6-diphosphate (FDP)] had significantly higher contribution scores than non-phosphorylated substrates (glucose and pyruvate). This finding aligns with the glycolysis process, where phosphorylated substrates have higher binding specificity to enzyme molecules, thereby promoting the forward reaction. Based on these prediction results, we decided to retain spermidine and folinic acid while discarding the other transcription-translation factors. For energy substrates, we selected PEP, which has the highest contribution score of 1.4.

### 2.4. Concentration optimization of the simplified CFE formulation

Having established a simplified composition, we proceeded to the second screening round to optimize the concentrations of the selected additives. We used 3 fluorescent colors to encode 5^3^ (125) combinations of PEP, spermidine, and folinic acid concentrations (Fig. 4a). The in-droplet screening produced 61,305 datasets (Fig. 4b), which were subsequently binned into the 125 combinations. For in silico optimization, we chose extreme gradient boosting (XGBoost, Fig. 4c), a scalable decision tree-based strategy widely used for enzyme activity prediction and drug discovery. The XGBoost model was initiated with a training set of 100 experimental combinations. During the training process, the training set was divided into 5 subsets for cross-validation. The trained model was verified using the remaining 25 experimental combinations, exhibiting an R^2^ of approximately 0.99 and a linear fit with an intercept of -0.002 and a slope of 1.004 (Supplementary Fig. 3d).

**Figure 4.**
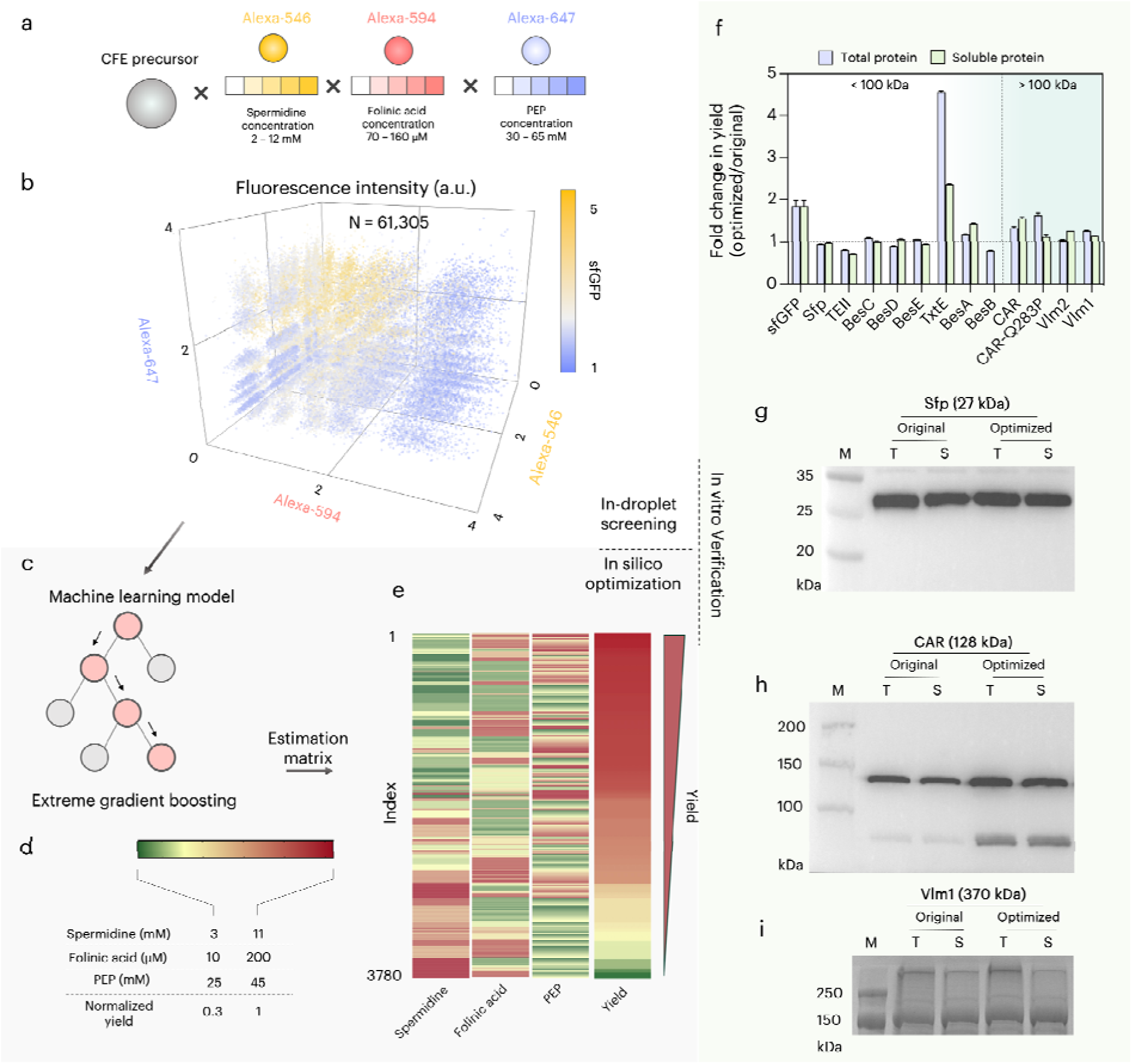
Second-round optimization of the *E. coli* extract-based CFE system. (a&b) In-droplet screening to optimize the concentrations of the selected components. (a) The fluorescence coding strategy. (b) A 3D scatter plot showing the fluorescence intensity profile of the droplets. The color of the dots indicates the sfGFP intensity. (c-e) In silico optimization. (c) An extreme gradient boosting model used in the optimization. (d) Concentration ranges scanned in the model prediction. (e) The estimation matrix sorted by yield level. (f-i) In vitro verification. (f) Fold change in the expression yield of 12 proteins with the simplified and optimized CFE formulation compared to the original recipe. The error bars show technical errors in analyzing the gel electrophoresis images (N = 3). (g-i) Western plot results of (g) Sfp (27 kDa), (h) CAR (128 kDa), and (i) Vlm1 (370 kDa).

After validating the model, we used it to span a broader range of concentrations (Fig. 4d) and predict the yields of 3,780 combinations (Fig. 4e). The prediction revealed the highest yield at 40 mM PEP, 130 μM spermidine, and 4 mM folinic acid. To validate the prediction, we employed the highest-yield formula to express sfGFP in bulk and found that, compared to the original formula, the yield increased by 1.9-fold, from 0.71 to 1.38 mg/mL (Supplementary Fig. 4a). Meanwhile, the total cost of additives was reduced by 2.1-fold, from 1.87 to 0.90 USD per mL of CFE assay (Supplementary Table 1). As a result, the unit cost of expressing sfGFP with the CFE was reduced by 4-fold, from 2.63 to 0.72 USD/mg.

To further test the general applicability of the optimized formulation, we expressed 12 proteins of various molecular weights (27 to 370 kDa) in vitro (Supplementary Fig. 4b). The fold changes in expression levels compared to those obtained with the original formulation are detailed in Fig. 4f. As shown, the expression levels of five proteins, including Sfp (27 kDa, Fig. 4e), BesC (29 kDa), BesD (29 kDa), and BesE (29 kDa), were maintained. Notably, TxtE (45 kDa) experienced a 4.5-fold increase in total expression and a 2.3-fold increase in soluble expression. Additionally, BesA (50 kDa), CAR (128 kDa, Fig. 4f), CAR-Q283P (128 kDa), Vlm2 (284 kDa), and Vlm1 (370 kDa, Fig. 4g) showed noticeable increases in total and/or soluble expression. The expression of TEII (27 kDa) and BesB (54 kDa) was slightly lowered. Taken together, our streamlined protocol demonstrates comparable or enhanced expression of various proteins while markedly reducing the costs associated with CFE reactions. These advancements are crucial for leveraging CFE systems in the rapid, economical, and efficient production of therapeutics, chemicals, and materials.

### 2.5. Optimization of *B. subtilis*-based CFE by transfer learning

We next questioned if we could adapt the established *E. coli* model to fit a different chassis cell, thereby reducing the effort required to optimize CFE systems across various hosts. To achieve this, we employed transfer learning, a machine-learning approach that repurposes a pre-trained model for a different task by learning from a limited set of new data. To examine this concept, we selected *B. subtilis*, a Gram-positive bacterium widely used as a microbial cell factory for recombinant protein production^23^, as the new chassis cell (Fig. 5a). For the in-droplet screening, we started from the primary screening results from the *E. coli* system and generated 125 experimental combinations (N = 59,741) by varying the concentrations of spermidine, folinic acid, and PEP (Supplementary Fig. 5a). To evaluate the performance of transfer learning, we varied the experimental set size between 27 (3×3×3) and 125 (5×5×5) and compared the prediction accuracies with a model built directly from the learning set. As shown in Fig. 5b, the transfer learning approach exhibited high prediction accuracy (R^2^ > 0.95) even with a limited experimental scale (e.g., 27 and 36 combinations). In contrast, the direct learning approach required at least 75 experimental sets to achieve reasonable accuracy. For example, with a transfer model built on 36 combinations (3×4×3), we obtained an R^2^ of 0.99 and a linear fit with an intercept of -0.008 and a slope of 1.01, whereas the direct learning model yielded an R^2^ of 0.83, an intercept of - 0.49, and a slope of 1.50 (Fig. 5c). These results confirm the effectiveness of transfer learning in establishing new CFE models with minimal input from experimental scales.

**Figure 5.**
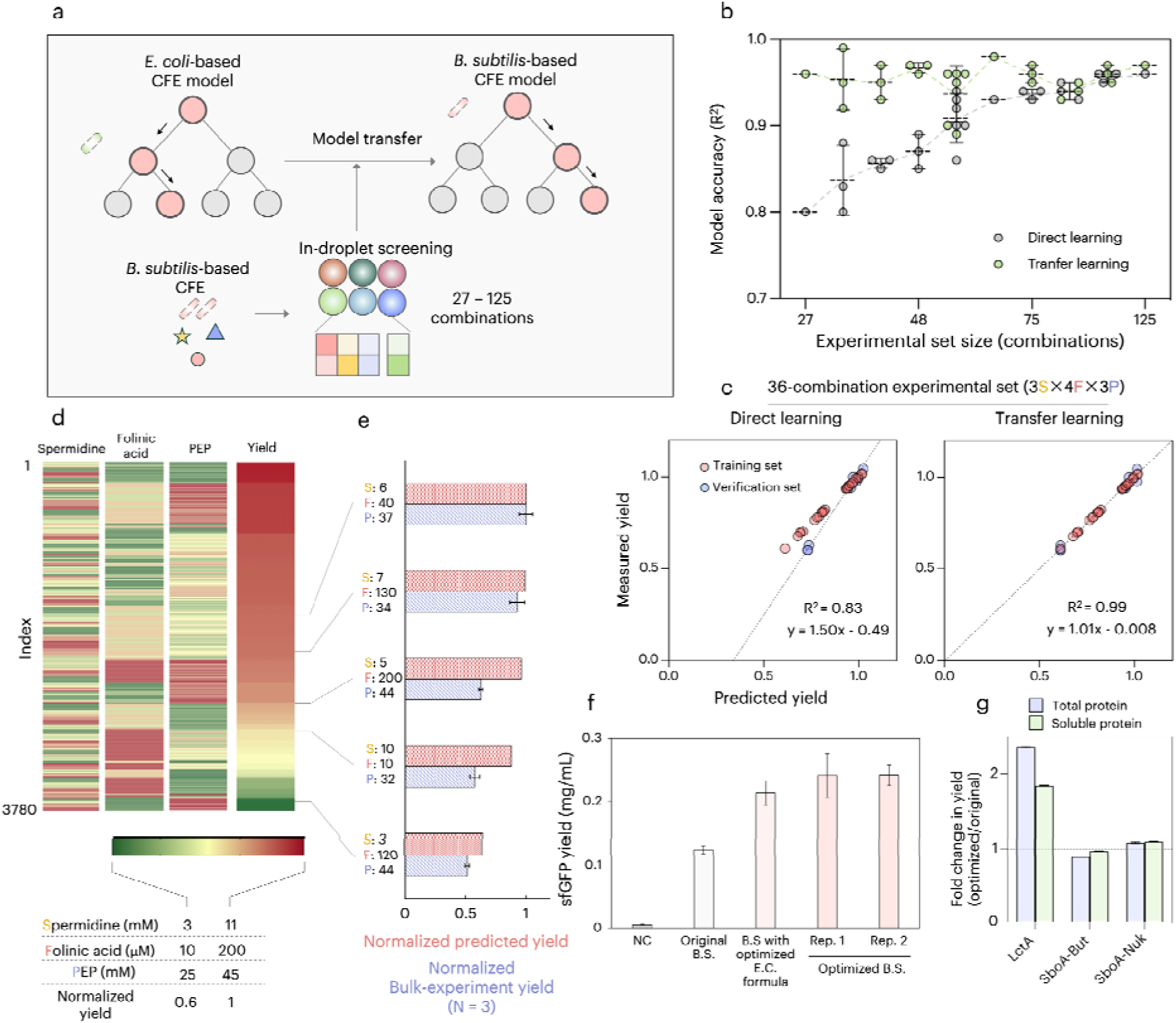
Optimization of a *B. subtilis* extract-based CFE system by transfer learning. (a) Schematic of the transfer learning process. The process initiates with the established *E. coli*-based model. By learning limited *B. subtilis* experimental data, the *E. coli* model is transferred to a *B. subtilis* model through a feature enhancement approach. (b) Model accuracy obtained at varying learning set sizes. The transfer learning approach is compared with a model directly built on the *B. subtilis* experimental data. (c) The measured sfGFP yield (in-droplet screening) vs. the predicted yield of a direct learning model and the transfer learning model. Both the models study 36 (3×4×3) experimental combinations. (d) The estimation matrix sorted by yield level. (e) Bulk sfGFP expression (N = 3) under 5 notified compositions and the model prediction results. (f) Experimental estimation of the optimized formulation to express sfGFP (N = 6). (g) Fold change in the expression yield of 3 peptides with the simplified and optimized CFE formulation compared to the original recipe. The error bars show technical errors in analyzing the gel electrophoresis images (N = 3).

Next, we utilized a 36-combination transfer model (3×4×3, R^2^ ∼ 0.99) to predict the yield of 3,780 distinct combinations of varying concentrations of spermidine, folinic acid, and PEP (Fig. 5d). For verification, we selected 5 combinations and conducted bulk CFE to express sfGFP. The results (Fig. 5e) indicated a consistent trend in CFE yield between the predictions and bulk experiments, although the exact yield levels did not perfectly match. To further evaluate the optimization, we chose a predicted high-yield combination of 5 mM spermidine, 100 μM folinic acid, and 35 mM PEP to express sfGFP in bulk, resulting in a yield of ∼0.24 mg/mL. This represented an approximate 2-fold increase compared to our previous *B. subtilis* recipe. Compared to *B. subtilis* CFE using the optimized *E. coli* formulation (the high-yield combination before model transfer, Fig. 4e), the yield further increased by 14% (Fig. 5f). Beyond sfGFP, the optimized and simplified formulation also demonstrated maintained or increased yields when expressing three short antimicrobial peptides (Fig. 5g and Supplementary Fig. 6). These results clearly demonstrate the effectiveness of DropAI in efficiently optimizing various CFE systems.

## 3. Discussion

CFE systems, particularly those based on cell lysates, represent promising platforms for various synthetic biology applications, including therapeutic development^8^, genetic part/circuit prototyping biomanufacturing^7,24^, artificial cell construction^25,26^, natural product biosynthesis^27-29^, and biomanufacturing^30,31^. Central to these applications is the need for consistent and efficient protein expression. In CFE reactions, supplemented components such as amino acids, nucleotides, salts, polyamines, and energy sources are essential alongside cell lysates and gene templates to ensure high-yielding protein production^1^. However, the cost of these additives can be prohibitive, driving up the overall expense of CFE systems. Therefore, a strong motivation exists to simplify CFE formulations to enhance productivity, reproducibility, ease of use, and reduce costs. Through DropAI optimization, we successfully streamlined the CFE formulation by eliminating five additives while maintaining or even enhancing protein expression levels compared to unoptimized reactions. Our findings highlight significant opportunities for improving protein expression in cell lysate-based CFE systems. Moreover, we demonstrated the transferability of our optimized recipe to another CFE system derived from *B. subtilis* using transfer learning. These results underscore the effectiveness and robustness of employing transfer learning to optimize novel CFE systems with minimal experimentation.

As this work has showcased, DropAI is an efficient, rapid, and cost-effective strategy for high-throughput screening (HTS) of CFE systems. During the two-round optimization of the *E. coli*-based CFE system, approximately 100,000 droplets were measured (Figs. 3f & 4b), corresponding to a total volume of only 25 μL. Considering the liquid lost during microfluidic processes, collection, transfer, and imaging, the overall reagent consumption is around 100 μL. The liquid handling in DropAI is fully streamlined with microfluidics, allowing combinatorial reactions to be prepared at 300 Hz. This means only 6 minutes are required to prepare 100,000 independent screening reactions. The overall operation time of DropAI is around 4 hours per round (15 min reagent preparation, 20 min microfluidic setup, 2 h satellite droplet generation, 25 min merging, and 1 h imaging). If the component-coding droplets are prepared in advance as stock, which is entirely feasible, the operation time can be reduced to approximately 2 hours. In contrast, screening an equivalent scale of reactions using well-plates and liquid-handling robot-based HTS platforms (with a unit reaction volume of around 10 μL) would consume over 1 liter of reagents, and the logistics of liquid handling could take several days^15,32,33^.

DropAI is a scalable platform. There is further room for improving DropAI’s throughput and combinatorial coding capacity. The microfluidics used in this work include droplet generation and multi-droplet merging. These techniques can be parallelized to enhance throughput^34-36^, allowing approximately 10^8^ combinatorial reactions to be conducted per day. Additionally, the combinatorial coding capacity can be expanded by incorporating more fluorescent colors and engineering the microfluidics to support the grouping and merging of more droplets. For instance, using a commercial 6-color droplet reader, such as the BioRad QX600, the combinatorial coding capacity could reach a scale of 10^6^. This high scalability makes DropAI a versatile platform that combines high-throughput experimentation with artificial intelligence. The combinatorial screening of CFE systems presented here uses only part of its capacity. DropAI may also be applied to various other scenarios, such as combinatorial drug screening^37-39^, chemical process optimization^40^, and catalyst discovery^41^. This versatility and efficiency make DropAI a powerful tool for advancing research and development in synthetic biology and beyond.

## 4. Methods

### 4.1. Microfluidic device fabrication

The microfluidic devices (Supplementary Fig. 1a) were designed using AutoCAD and printed as dark-field plastic photomasks. To fabricate the devices, a 3-inch wafer was coated with photoresist SU-8 3025 (Microchem) using a spin coater in a two-step process: 30 seconds at 500 rpm, followed by 60 seconds at 2500 rpm. The wafer was soft-baked on a hotplate at 95°C for 20 minutes, exposed to a 120 mW lamp (M365L2, Thorlabs) for 3 minutes and 20 seconds under a photomask, and baked again at 95°C for 4 minutes. The repeated process was applied if a second layer was needed. Next, the wafer was incubated in SU-8 developer (Microchem) for 10 minutes, and excess developer was removed with isopropanol and ethanol, followed by blow-drying with a nitrogen gun. The wafer was then placed in a plastic petri dish, covered with a PDMS precursor (SYLGARD 184, Dow Corning) mixed with a curing agent in a 10:1 (w/w) ratio, degassed in a vacuum chamber, and cured at 60°C for 8 hours. The cured PDMS slab with the pattern was peeled off from the mold, and inlet/outlet ports were created using a 0.7 mm hole puncher. The patterned PDMS slab was bonded to a glass slide using oxygen plasma treatment. Before use, the devices were treated with Aquapel (PPG Industries) to render the channel surfaces hydrophobic.

### 4.2. Strains, media, and plasmids

*E. coli* DH5α was used for coloning and plasmid propagation. *E. coli* BL21 Star (DE3) and *B. subtilis* 164T7P were employed for cell extract preparation, respectively. LB medium (10 g/L sodium chloride, 5 g/L yeast extract, and 10 g/L tryptone) was used for cultivating bacterial cells. For cell extract preparation, cells were grown in 2×YTPG medium, which consists of 10 g/L yeast extract, 16 g/L tryptone, 5 g/L sodium chloride, 7 g/L potassium hydrogen phosphate, 3 g/L potassium dihydrogen phosphate, and 18 g/L glucose, adjusted to pH 7.2. The genes and plasmids used in this work are detailed in Supplementary Table 2.

### 4.3. Preparation of cell extracts

Cell extracts were prepared as previously described^18,22^. Briefly, 1 L of 2×YTPG medium was inoculated with an overnight culture to an initial OD600 of 0.05. When the OD600 reached 0.6-0.8, cells were induced with 1 mM IPTG and harvested at an OD600 of 3.0. The cells were washed three times with cold S30 Buffer (10 mM Tris-acetate, 14 mM magnesium acetate, and 60 mM potassium acetate). The cell pellet was resuspended in S30 Buffer (1 mL/g of wet cell mass) and lysed by sonication (10 s on/off, 50% amplitude, input energy ∼600 Joules). The lysate was centrifuged twice at 12,000 g for 10 minutes at 4°C. The resulting supernatant was flash-frozen in liquid nitrogen and stored at -80°C until use.

### 4.4. Cell-free gene expression

The CFE reactions were performed in 1.5 mL centrifuge tubes (15 μL) or 80-μm droplets (250 pL) at 30°C for 4 hours. Each reaction contained 27% (v/v) cell extract, 13.3 μg/mL plasmid, 33 mM energy substrate (PEP, FDP, F6P, glucose, or pyruvate), and 35 other components. The component information for the original and optimized formulations of *E. coli* and *B. subtilis* CFE systems is detailed in Supplementary Table 1. The droplet CFE formulation contained additional 0.2% (v/v) P-188 and 1% (v/v) PEG-6000 to stabilize the emulsions. Synthesized proteins were analyzed using SDS-PAGE and Western blotting. The concentration of sfGFP was determined based on fluorescence intensity measured with a microplate reader (SYNETGY H1). For this, 2 μL of the CFE reaction was diluted with 48 μL nuclease-free water and placed in a flat-bottom 96-well plate. Measurements of sfGFP fluorescence were taken with excitation at 485 nm and emission at 528 nm. The fluorescence was converted to concentration (mg/mL) using a linear standard curve created in-house. For other proteins, their Western blot band densities and areas were analyzed by ImageJ, which were used to estimate the fold changes in yield between the optimized and original recipes. For in-droplet screening, the CFE mix was prepared at 1.3-fold of the final concentration, considering dilution in the merging process. The screening components were also removed from the CFE mix and prepared in the satellite droplets.

### 4.5. Western blot

The CFE reaction mixture was centrifuged at 12,000g for 10 minutes at 4°C, and the supernatant was collected as the soluble fraction. Both 10 μL of total protein and 10 μL of soluble protein were mixed with 10 μL of 2x loading buffer and heated at 98°C for 10 minutes. Each sample (10 μL) was then loaded onto an SDS-PAGE gel and transferred to a PVDF membrane (Bio-Rad) using 1x transfer buffer (25 mM Tris-HCl, 192 mM glycine, 20% methanol, pH 8.3). The membrane was blocked with Protein Free Rapid Blocking Buffer (EpiZyme) for 30 minutes at room temperature, followed by three washes with 1x TBST buffer (10 mM Tris-HCl, 150 mM NaCl, 0.1% Tween 20, pH 7.5). It was then incubated with His-Tag Mouse Monoclonal Antibody (Proteintech) diluted 1:10000 in TBST buffer for 1 hour, washed three more times with 1x TBST buffer, and incubated with HRP-Goat Anti-Mouse IgG (H+L) Antibody (Proteintech) diluted 1:10000 in TBST buffer for another hour. After a final set of three washes, the membrane was visualized using Omni ECL reagent (EpiZyme) under UVP ChemStudio (analytikjena).

### 4.6. Microfluidic operations

The microfluidic experiments were conducted at a custom microfluidic station consisting of an inverted microscope equipped with a fast-speed camera and 6 syringe pumps. The oil [HFE-7500 (3M) and 2.5% wt PEG-PFPE surfactant (Dapu Biotechnology)] and prepared aqueous solutions were stored in 1-mL plastic syringes and connected to the microfluidic chip mounted on the microscope through plastic tubing (SCI PE/2). The flowrates applied in each microfluidic experiment are detailed in Supplementary Table 3. For droplet merging, a 1.5 kV (Vpp), 50 kHz AC voltage was applied to the electrode port using an inverter (TDK CXA-L0605-VJL) circuit.

### 4.7. In-droplet screening

The screening components, their concentrations, fluorescent dye, and dye concentrations in proof-of-concept experiments and each screening round are detailed in Supplementary Table 4. The collected droplets were loaded onto a cell counting slide (Thermal Fisher) for microscopic observation (Nikon, Eclipse Ti2) and to a 4-channel digital PCR droplet reader (Nebula Reader, Dapu Biotechnology) for multi-channel imaging. The fluorescent intensities of the droplets were obtained with a built-in droplet analysis software (Nebula-Astrolabe, Supplementary Fig. 1d). For the clustering of 9^4^ combinations, the raw droplet intensity data was filtered to eliminate marginal points with a multi-band-pass filter program built in MATLAB. For other experiments, the raw data was used without filtering. The data was categorized into different combination bins with a custom MATLAB program.

### 4.8. AI models

We used the neural network model^42,43^ to evaluate the contribution of the 12 additives to sfGFP yield. The data used to train the combinatorial screening model (CSM) was obtained directly from the in-droplet screening. The droplet experiment tested 100 combinations, each with more than 300 repeats. The experimental data was processed before training. The sfGFP yield (fluorescent intensity) was normalized to the maximum value among the data. The normalized yield of each combination was averaged, and the mean values were used. In each combination, we used one-hot encoding to describe the component’s existence, with “1” indicating the presence and “0” indicating the absence (Supplementary Fig. 3a). Among the 100 sets of experimental results, 70 are for training and 30 are for validation. The established model traversed the entire combinatorial space and predicted the yield of each combination. Note that every combination was set to have no more than one energy substrate. Thus, the entire combinatorial space contained 768 [2^7^×(5+1)] combinations. After prediction, we evaluated the contribution of each additive to the yield by comparing the yields with a certain additive present to the yields with it absent (Supplementary Fig. 3c). CSM is a fully connected model compiled based on Python (version 3.10.11) and TensorFlow (version 2.8.0). It consists of 7 fully connected layers. The training is compiled with the Adam optimizer.

In the second round of optimization, we used the XGBoost algorithm^44^ to train the concentration model. The in-droplet screening generated 125 combinations of varying concentrations of spermidine, folinic acid, and PEP. Among the 125 combinations, 100 combinations were randomly selected as the training set and the remaining 25 as the validation set. The input of the model was the concentration of the three components, and the output was the sfGFP yield (fluorescence intensity). The learning set was first standardized with ‘StandardScaler’ in Python’s scikit-learn library, converting the distributions to have a mean of 0 and a variance of 1. Then, the learning started with hyperparameter tuning with the XGBoost regression model using GridSearchCV. A 5-fold cross-validation was employed, and model performance was evaluated using negative mean squared error (neg_mean_squared_error). The model with the best hyperparameters was finally selected, and its accuracy was estimated using the validation sets. The verified model was used to predict 3,840 combinations and the yield was normalized to the largest value in the prediction (Fig. 4e). The model was compiled in Python (version 3.10.11).

### 4.9. Transfer learning

To transfer the *E. coli* model to fit *B. subtilis-*based CFE, we employed the following steps: 1) Original Model Prediction: we used the trained *E. coli* model to predict the standardized training and verification sets data in *B. subtilis*-based CFE system, obtaining prediction values for both sets. 2) Feature Enhancement: We added the prediction values as new features to the original feature set of *B. subtilis*-based CFE, forming an enhanced feature set. 3) New Model Training: Using this enhanced feature set, a new semi-quantitative prediction model (again using XGBoost) was trained for the yield levels in the B. subtilis extract-based CFE system. To evaluate the minimal experimental input for transfer learning, the 125 experimental sets were down-scaled to 27 different subsets (Fig. 5b and Supplementary Table 4). 80% of the experimental data was used for training and 20% for verification. A new transferred model was obtained from each of the subsets, and their coefficient of determination (R^2^) was determined (Supplementary Fig. 5b). For comparison, we also built a model directly trained from each of these experimental subsets. Here, the training process was identical to that of the *E. coli* model. Out of 27 transfer-learning models, one (Fig. 5c) was selected to predict 3,840 combinations and the yield was normalized to the largest value in the prediction.

