## Supplementary Information for "AI-driven high-throughput droplet screening of cell-free gene expression"

**Supplementary Table 1.** Formulation and cost of CFE systems used in this work.
Please refer to the supplementary .xlsx file.

**Supplementary Table 2.** Genes and plasmids used in this work.
Please refer to the supplementary .xlsx file.

**Supplementary Table 3.** Flowrates applied in the microfluidic devices.

| Device | Flowrate (µL/h) |
|---|---|
| Satellite droplet generation<br>(Supplementary Fig. 1) | Oil: 600<br>Reagent: 300 |
| Carrier droplet generation | Oil: 1000<br>CFE mix: 500 |
| Drop grouping and merging | Oil: 800<br>CFE drop: 300<br>Satellite drop: 40<br>Spacer oil: 100 |
| Drop grouping and merging<br>(simplified) | Oil: 800<br>CFE mix: 300<br>Satellite drop: 40<br>Spacer oil: 100 |

**Supplementary Table 4.** Fluorescent coding conditions in the droplet experiments.

4.1. Dyes and their concentrations used in combinatorial library characterization (Fig. 2)

| Dye | Alexa-488 (nM) | Alexa-546 (nM) | Alexa-594 (nM) | Alexa-647 (nM) |
|---|---|---|---|---|
| Concentration 1 | 0 | 0 | 0 | 0 |
| Concentration 2 | 100 | 100 | 100 | 100 |
| Concentration 3 | 200 | 200 | 200 | 200 |
| Concentration 4 | 300 | 300 | 300 | 300 |
| Concentration 5 | 400 | 400 | 400 | 400 |
| Concentration 6 | 500 | 550 | 550 | 500 |
| Concentration 7 | 650 | 700 | 700 | 650 |
| Concentration 8 | 800 | 900 | 900 | 800 |
| Concentration 9 | 1000 | 1200 | 1200 | 1000 |

4.2. Alexa-647 and the corresponding $Mg^{2+}$ concentrations used in Fig. 3b.

| Alexa-647 concentration (nM) | $Mg^{2+}$ concentration (mM) |
|---|---|
| 300 | 0 |
| 900 | 12 |

4.3. Alexa-647 and the corresponding $Mg^{2+}$ concentrations used in Fig. 3c.

| Alexa-647 concentration (nM) | $Mg^{2+}$ concentration (mM) |
|---|---|
| 0 | 24 |
| 300 | 20 |
| 700 | 16 |
| 900 | 12 |
| 1600 | 8 |

4.4. Coding conditions in the first round of in-droplet screening of the *E.coli* CFE (Fig. 3d).

| Dye | Concentration (nm) | Encoded condition |
|---|---|---|
| Alexa-546 | 84 | $K_2CO_3$ + NAD + CoA |
| Alexa-546 | 200 | NAD + CoA |
| Alexa-546 | 330 | $K_2CO_3$ + CoA |
| Alexa-546 | 500 | $K_2CO_3$ + NAD |
| Alexa-594 | 160 | Folinic acid + spermidine + putrescine + tRNA |
| Alexa-594 | 330 | Spermidine + putrescine + tRNA |
| Alexa-594 | 500 | Folinic acid + putrescine + tRNA |
| Alexa-594 | 830 | Folinic acid + spermidine + tRNA |
| Alexa-594 | 1200 | Folinic acid + spermidine + putrescine |
| Alexa 647 | 160 | PEP |
| Alexa 647 | 330 | F6P |
| Alexa 647 | 500 | FDP |
| Alexa 647 | 830 | Pyr |
| Alexa 647 | 1200 | Glu |

4.5. Coding conditions in the second round of in-droplet screening of the *E.coli* CFE (Fig. 4).

| Dye | Concentration (nm) | Encoded condition |
| --- | --- | --- |
| Alexa-546 | 84 | Spermidine (2.8 mM) |
| Alexa-546 | 200 | Spermidine (4.1 mM) |
| Alexa-546 | 330 | Spermidine (6.7 mM) |
| Alexa-546 | 500 | Spermidine (9.3 mM) |
| Alexa-546 | 700 | Spermidine (11.9 mM) |
| Alexa-594 | 160 | Folinic acid (70 μM) |
| Alexa-594 | 330 | Folinic acid (100 μM) |
| Alexa-594 | 500 | Folinic acid (120 μM) |
| Alexa-594 | 830 | Folinic acid (140 μM) |
| Alexa-594 | 1200 | Folinic acid (160 μM) |
| Alexa 647 | 160 | PEP (31 mM) |
| Alexa 647 | 330 | PEP (40 mM) |
| Alexa 647 | 500 | PEP (48 mM) |
| Alexa 647 | 830 | PEP (56 mM) |
| Alexa 647 | 1200 | PEP (65 mM) |

4.6. Coding conditions in the in-droplet screening of the *B.subtilis* CFE (Fig. 5).

| Dye | Concentration (nm) | Encoded condition | Downscale to 3 levels | Downscale to 4 levels |
|---|---|---|---|---|
| Alexa-546 | 84 | Spermidine (3 mM) | ✓ | ✓ |
| Alexa-546 | 200 | Spermidine (5 mM) | | |
| Alexa-546 | 330 | Spermidine (7 mM) | ✓ | ✓ |
| Alexa-546 | 500 | Spermidine (9 mM) | | ✓ |
| Alexa-546 | 700 | Spermidine (11 mM) | ✓ | ✓ |
| Alexa-594 | 160 | Folinic acid (10 µM) | ✓ | ✓ |
| Alexa-594 | 330 | Folinic acid (50 µM) | | |
| Alexa-594 | 500 | Folinic acid (100 µM) | ✓ | ✓ |
| Alexa-594 | 830 | Folinic acid (150 µM) | | ✓ |
| Alexa-594 | 1200 | Folinic acid (200 µM) | ✓ | ✓ |
| Alexa 647 | 160 | PEP (25 mM) | ✓ | ✓ |
| Alexa 647 | 330 | PEP (30 mM) | | |
| Alexa 647 | 500 | PEP (35 mM) | ✓ | ✓ |
| Alexa 647 | 830 | PEP (40 mM) | | ✓ |
| Alexa 647 | 1200 | PEP (45 mM) | ✓ | ✓ |

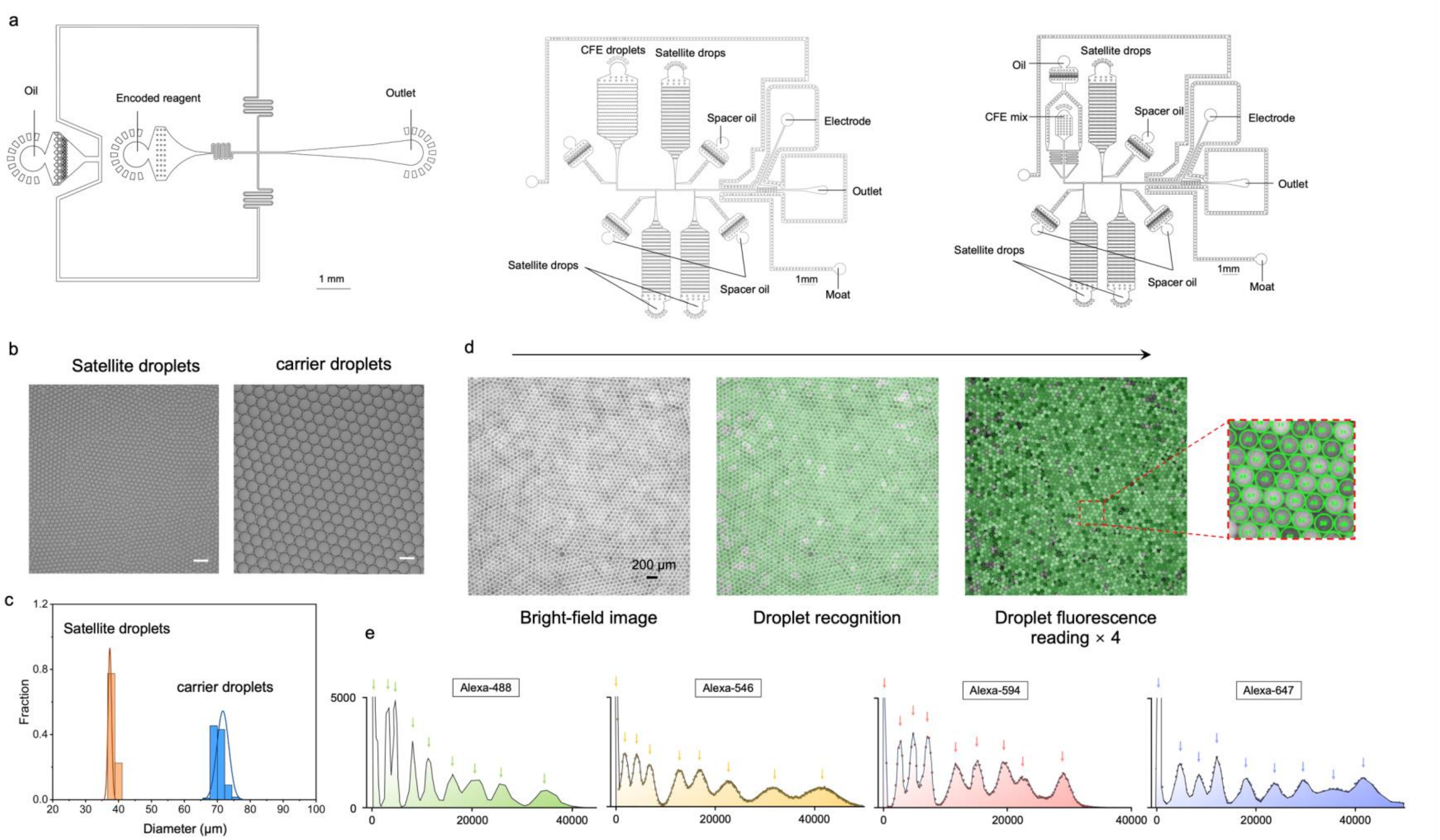


**Supplementary Figure 1.** (a) The layout of microfluidic devices used in this work. (b) Representative micrographs depicting the satellite droplets and carrier droplets. Scale bars: 100 µm. (c) Diameter distribution of these droplets. (d) The four-channel droplet imaging workflow. (e) Droplet fluorescent distributions. The droplets (N = 206,634) encode 9 fluorescent levels in each color.

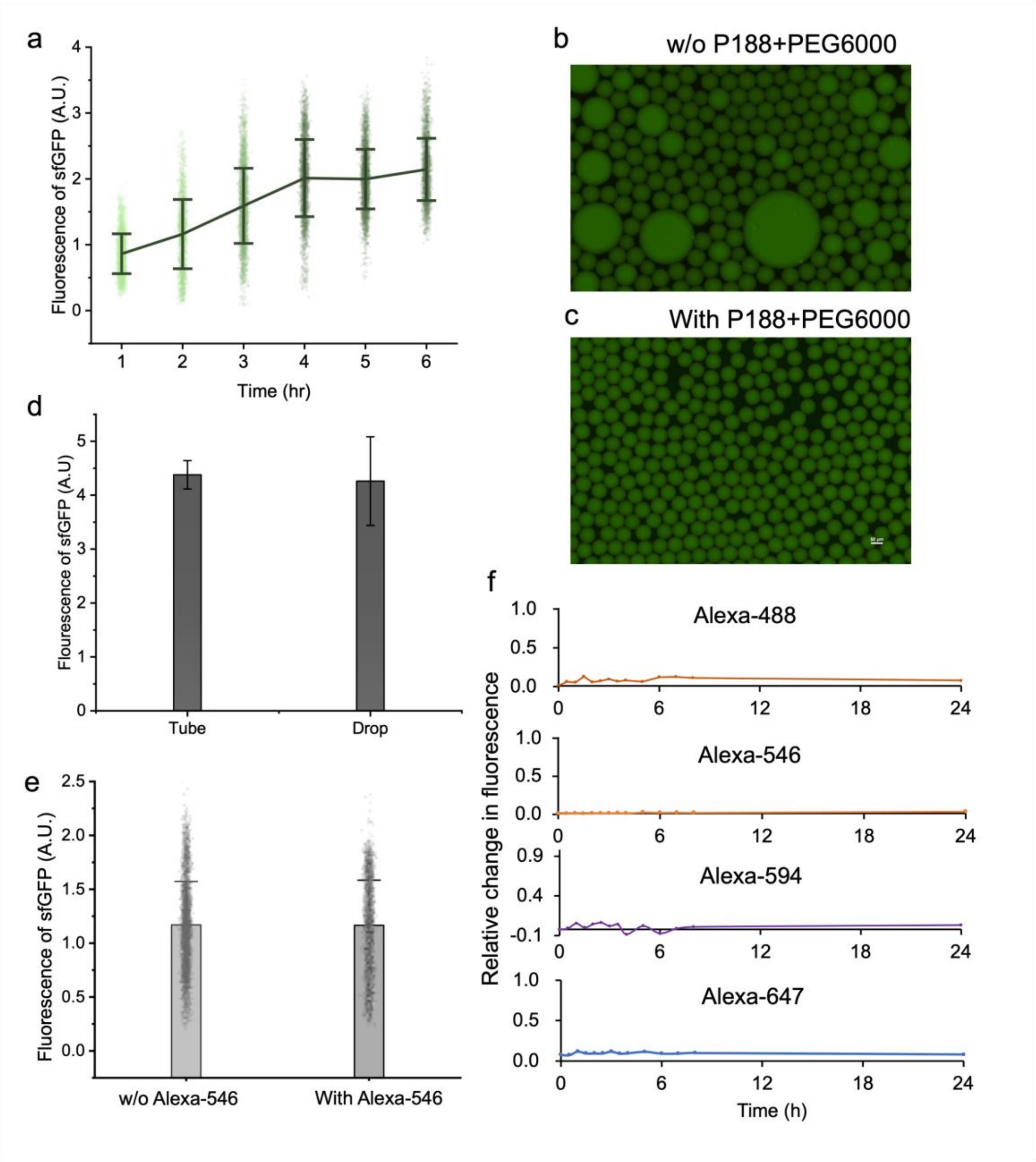


**Supplementary Figure 2.** (a) Fluorescent intensity of droplet-based CFE expressing sfGFP over time. (b&c) Representative fluorescent (sfGFP) micrographs depicting the effect of P188 and PEG6000 in stabilizing the droplets. Scale bar: 50 µm. (d) Comparison of the produced sfGFP intensity between the tube-based and droplet-based CFE systems. (e) sfGFP intensities in the CFE droplets with and without the fluorescent dye. (f) Change of droplet fluorescence (dye concentration: 100 nM) over time.

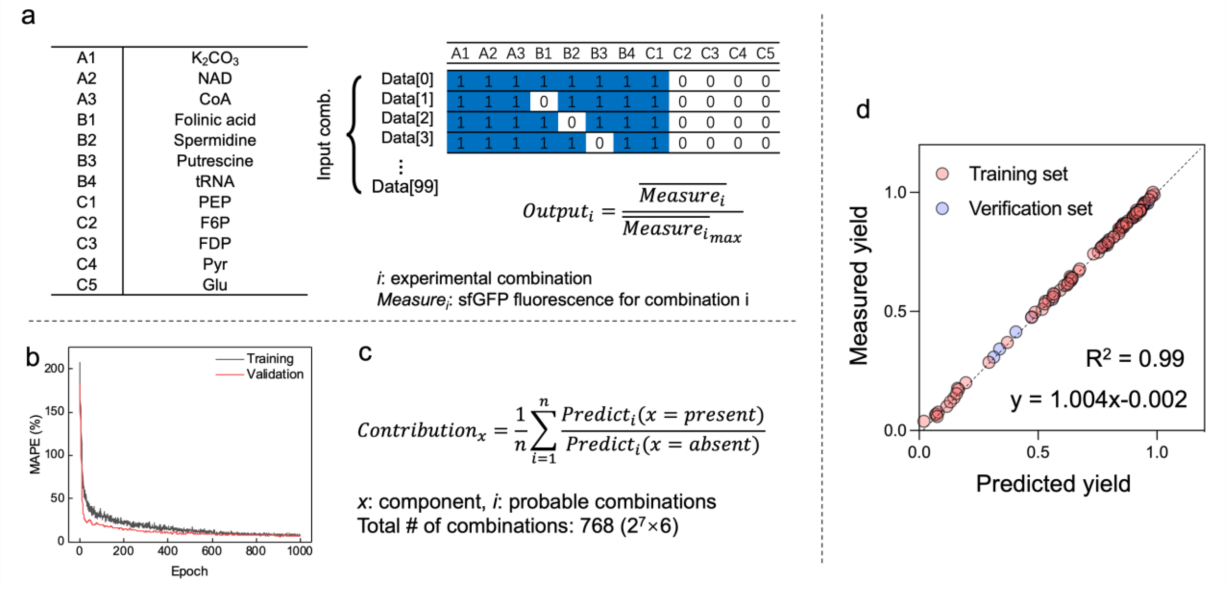


**Supplementary Figure 3.** (a) Component coding conditions and AI model input/output in the primary screening of an *E. coli*-based CFE system. (b) Mean absolute percentage error (MAPE) plotted as a function of training epochs. (c) Equation to evaluate the contribution of a component. (d) Model validation in the second-round optimization of the *E. coli*-based CFE system: sfGFP yield (fluorescence intensity) obtained from the droplets vs. the model predictions.

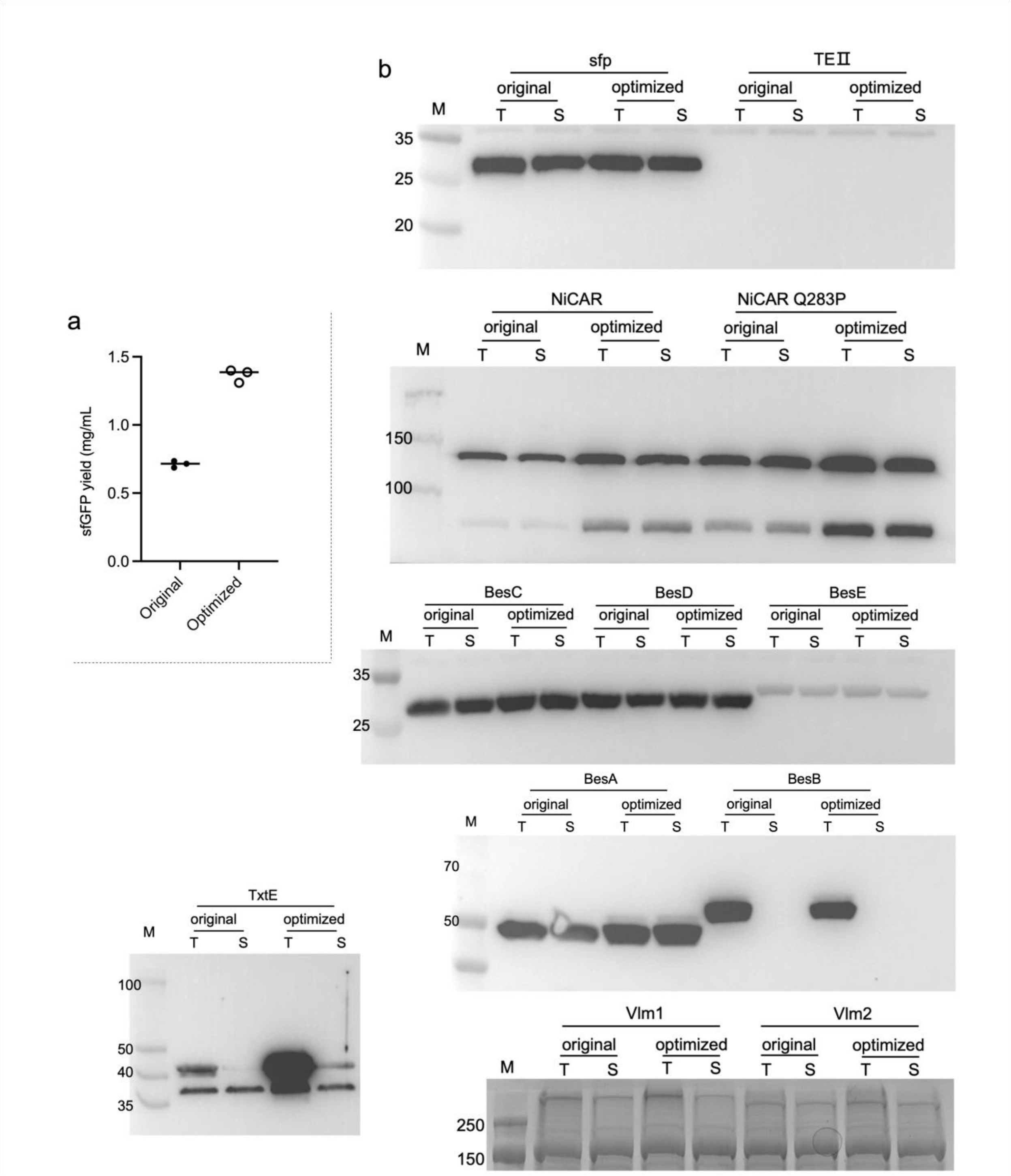


**Supplementary Figure 4.** (a) Bulk (15 µL) sfGFP yield of the original and optimized *E. coli*-based CFE systems. (b) Western blot results of 12 proteins expressed using the original and optimized formulation.

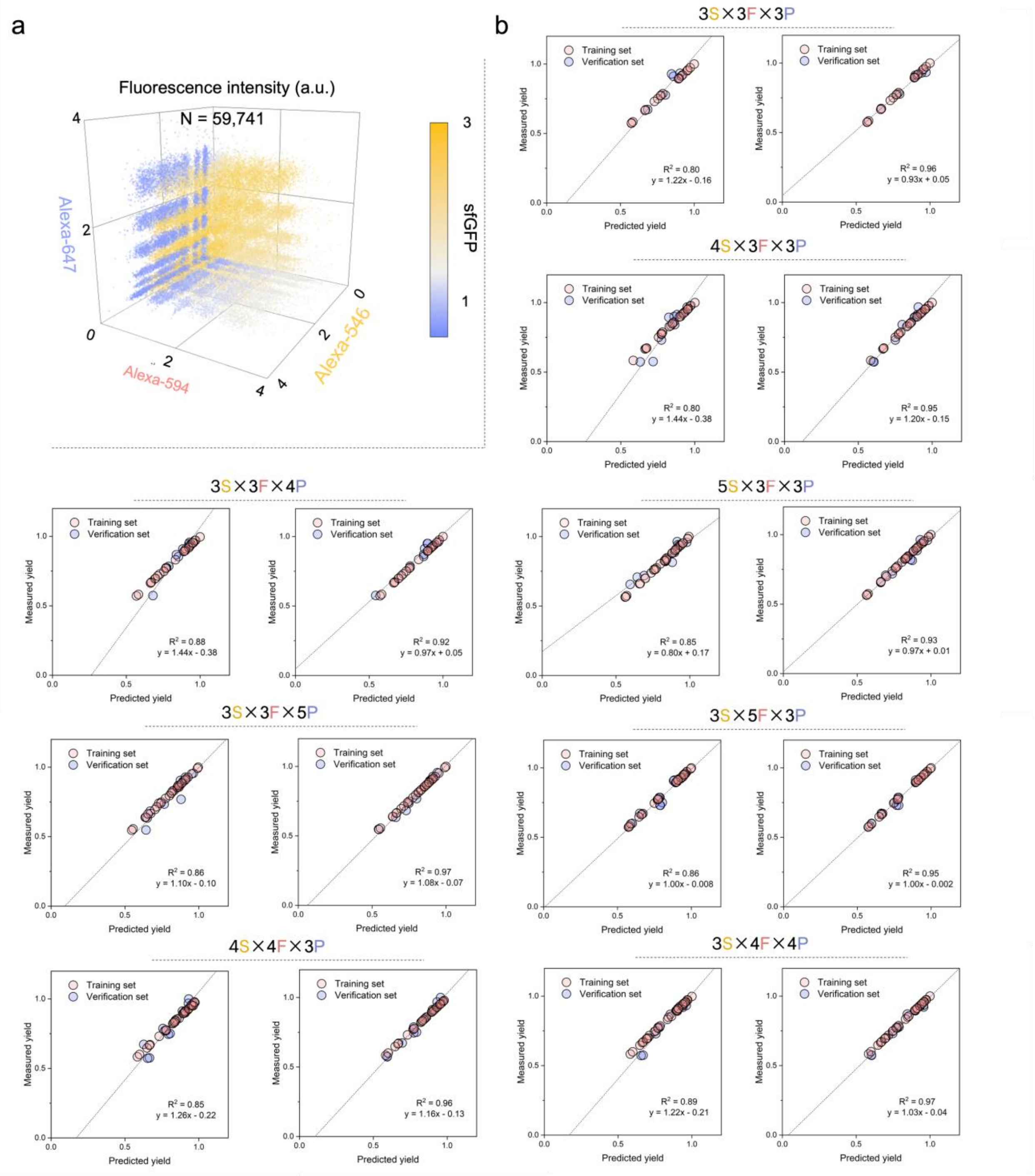

a
Fluorescence intensity (a.u.)
N = 59,741
Alexa-647
Alexa-594
Alexa-546
sfGFP
b
3S×3F×3P
Training set
Verification set
Measured yield
Predicted yield
R^2 = 0.80
y = 1.22x - 0.16
R^2 = 0.96
y = 0.93x + 0.05
4S×3F×3P
R^2 = 0.80
y = 1.44x - 0.38
R^2 = 0.95
y = 1.20x - 0.15
3S×3F×4P
R^2 = 0.88
y = 1.44x - 0.38
R^2 = 0.92
y = 0.97x + 0.05
5S×3F×3P
R^2 = 0.85
y = 0.80x + 0.17
R^2 = 0.93
y = 0.97x + 0.01
3S×3F×5P
R^2 = 0.86
y = 1.10x - 0.10
R^2 = 0.97
y = 1.08x - 0.07
3S×5F×3P
R^2 = 0.86
y = 1.00x - 0.008
R^2 = 0.95
y = 1.00x - 0.002
4S×4F×3P
R^2 = 0.85
y = 1.26x - 0.22
R^2 = 0.96
y = 1.16x - 0.13
3S×4F×4P
R^2 = 0.89
y = 1.22x - 0.21
R^2 = 0.97
y = 1.03x - 0.04

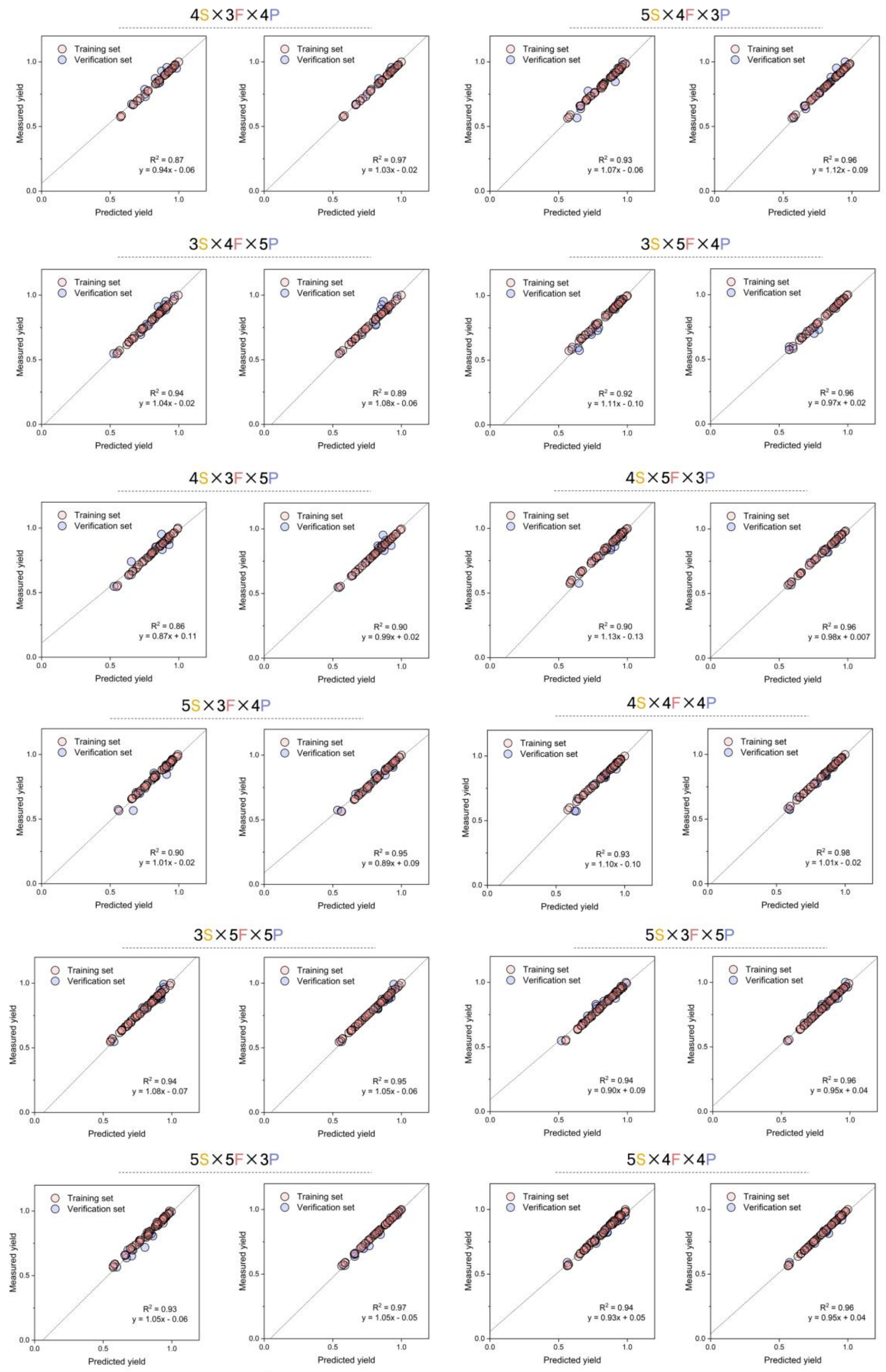
4S×3F×4P
Training set
Verification set
Measured yield
Predicted yield
R² = 0.87
y = 0.94x - 0.06
R² = 0.97
y = 1.03x - 0.02
5S×4F×3P
R² = 0.93
y = 1.07x - 0.06
R² = 0.96
y = 1.12x - 0.09
3S×4F×5P
R² = 0.94
y = 1.04x - 0.02
R² = 0.89
y = 1.08x - 0.06
3S×5F×4P
R² = 0.92
y = 1.11x - 0.10
R² = 0.96
y = 0.97x + 0.02
4S×3F×5P
R² = 0.86
y = 0.87x + 0.11
R² = 0.90
y = 0.99x + 0.02
4S×5F×3P
R² = 0.90
y = 1.13x - 0.13
R² = 0.96
y = 0.98x + 0.007
5S×3F×4P
R² = 0.90
y = 1.01x - 0.02
R² = 0.95
y = 0.89x + 0.09
4S×4F×4P
R² = 0.93
y = 1.10x - 0.10
R² = 0.98
y = 1.01x - 0.02
3S×5F×5P
R² = 0.94
y = 1.08x - 0.07
R² = 0.95
y = 1.05x - 0.06
5S×3F×5P
R² = 0.94
y = 0.90x + 0.09
R² = 0.96
y = 0.95x + 0.04
5S×5F×3P
R² = 0.93
y = 1.05x - 0.06
R² = 0.97
y = 1.05x - 0.05
5S×4F×4P
R² = 0.94
y = 0.93x + 0.05
R² = 0.96
y = 0.95x + 0.04

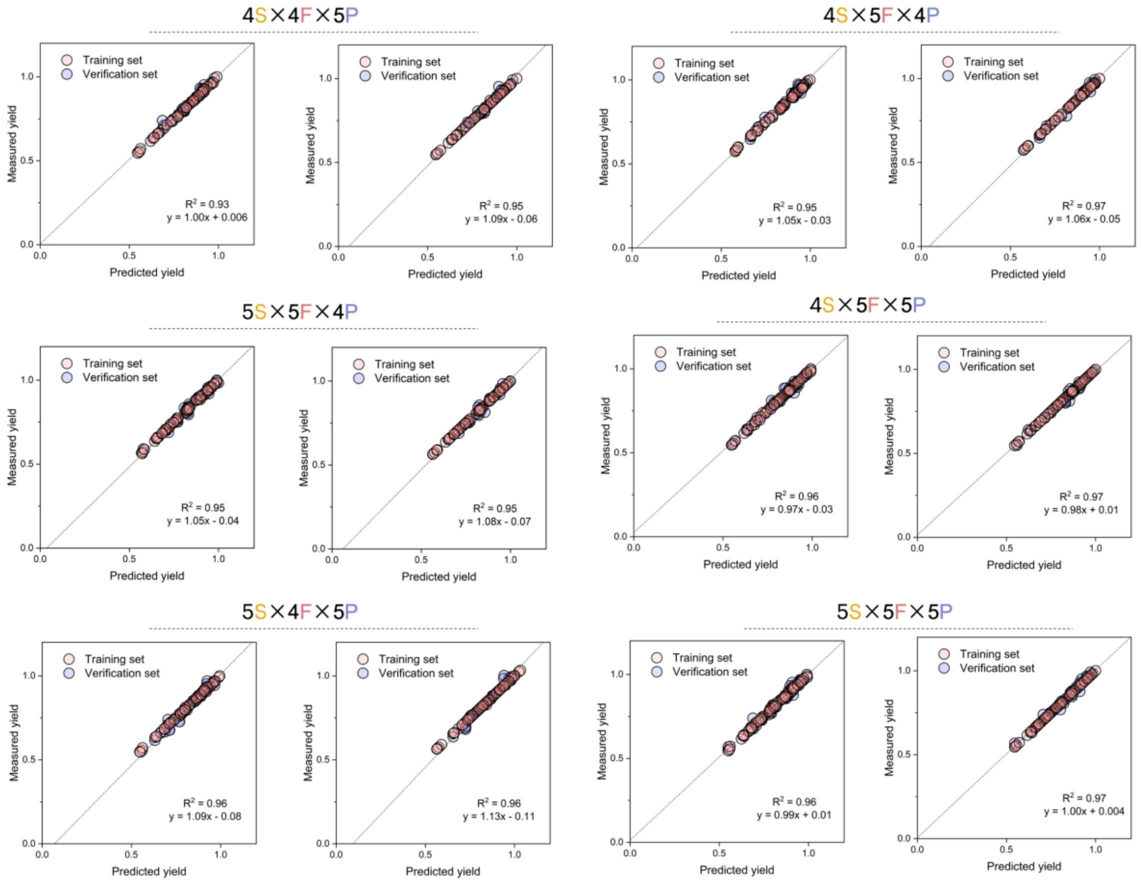


**Supplementary Figure 5.** (a) The fluorescence intensity profile of the droplets (N = 59,741) in the combinatorial screening of a *B. subtilis*-based CFE system. The color of the dots indicates the sfGFP intensity. (b) sfGFP yield (fluorescence intensity) obtained from the droplets vs. the model predictions of 26 groups of AI models. Each group studies a different subset of 125 experimental sets. The left panel in a group is the model directly built from the input data, while the right is the model transferred from the pre-established *E. coli* CFE model. The downscaled subsets of each additive are marked in Supplementary Table 4.6.

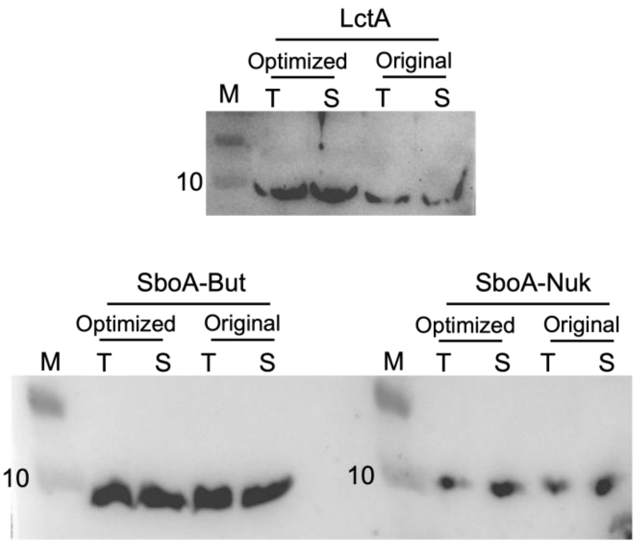


**Supplementary Figure 6.** Western blot results of 3 anti-microbial peptides expressed using the original and optimized *B. subtilis*-based CFE systems.
